# Pregnancy affects flight behavior in big brown bats (*Eptesicus fuscus*)

**DOI:** 10.1101/2024.01.08.574405

**Authors:** Adrienne Calistri-Yeh, Vanessa K Hilliard Young, Michaele Klingerman, Laura N. Kloepper

**Affiliations:** School for Environment and Sustainability, University of Michigan, Ann Arbor, MI, USA; Department of Biology, Saint Mary’s College, Notre Dame, IN, USA; Saint Joseph County Parks, South Bend, IN, USA; Department of Biological Sciences and Center for Acoustics Research and Education, University of New Hampshire, Durham, NH, USA

**Keywords:** bats, flight, pregnancy, wing-beat rate (WBR)

## Abstract

Pregnancy involves increased body mass and decreased locomotor performance in many species and can be especially impactful on volant animals. To test the hypothesis that bats modify flight behavior to adjust for pregnancy-related increases in mass, we recorded thermal video from a maternity colony of big brown bats (*Eptesicus fuscus*) as they emerged from a roost, during periods associated with pregnancy and post-pregnancy. From video tracking, we calculated the vertical drop distance before upward motion (UM), time from emergence until UM, average speeds before and after UM, and average wingbeat rate per second (WBR). Bats recorded during the pregnancy period had a significantly larger drop distance, longer time until UM, faster flight speed before UM, and higher WBR compared to bats recorded during the non-pregnancy period. Our results suggest that pregnancy has a significant effect on flight in female bats, with a particularly strong impact on achieving UM after emergence. However, the higher WBR recorded from bats flying during the pregnancy period implies that bats acclimate to such changes in body mass by altering their flight behaviors to sustain UM while pregnant.

## INTRODUCTION

Pregnancy involves morphological changes and increased body mass associated with decreased locomotor performance in terrestrial, aquatic, arboreal, and aerial species (Fleuren 2017; Smith and Young 2021); this reduces fitness by increasing predation risk and restricting ability to meet energetic demands (Fleuren 2017; Guillemette and Ouellet 2005; Hückstädt et al. 2018; Kullberg et al. 2002; Kullberg et al. 2005; Lee et al. 1996; Miles et al. 2000; Noren et al. 2011; Plaut 2002; Winfield and Townsend 1983). For example, gravid females demonstrate decreased locomotor speed and/or endurance compared to non-gravid females across a range of taxa (lizards: Miles et al. 2000, Shine 1980; snakes: Seigel et al. 1987; copepods: Winfield and Townsend 1983; scorpions: Shaffer and Formanowicz 1996; fish: Plaut 2002; dolphins: Noren et al. 2011; seals: Hückstädt et al. 2018; birds: Kullberg et al. 2002; Kullberg et al. 2005; Lee et al. 1996; Videler et al. 1988; bats: Hughes and Rayner 1991, 1993, McLean and Speakman 2000). For flying animals, gravidity has an even greater impact because increased mass also increases wing loading (Hayssen and Kunz 1996; Pennycuick et al. 1989), which is inherently disadvantageous to aerial locomotion. Gliding species compensate with larger body sizes (Fokidis and Risch 2008a; Shine et al. 1998), but actively flying species must generate lift to overcome the force of gravity (Pennycuick et al. 1989; Swartz et al. 2012; Vogel 2013). Thus, a different strategy is needed to overcome the increased wing loading and power required to sustain flight during pregnancy (Fokidis and Risch 2008b; Guillemette and Ouellet 2005; Hayssen and Kunz 1996; McLean and Speakman 2000).

One way to partially mitigate the impact of increased wing loading is to increase wingbeat frequency, thus generating more power (Fleuren 2017; Swartz et al. 2012). This strategy has been noted for some artificially loaded birds and bats, as well as pregnant bats in a small flight room, which increase wingbeat rate and decrease flight speed (Hughes and Rayner 1991, 1993; Pennycuick et al. 1989; Videler et al. 1988). However, biomechanical strategies for free-ranging species that drop from higher positions before achieving upward motion and thus face the additional challenge of overcoming vertical momentum, remain unknown. This initial dropping stage is an essential step of emergence; if the bat cannot generate enough lift to achieve upward motion during the vertical drop, it will be unable to successfully take flight. This can have implications for roost selection of maternity colonies, especially if flight performance differs between pregnant and non-pregnant bats.

This study aimed to explore the effect of pregnancy on flight for bats emerging and dropping from a natural roost. Specifically, we studied the flight behavior of big brown bats (*Eptesicus fuscus*), which experience a 15-20% increase in mass during pregnancy (Schmidly and Bradley 2016). We predicted that pregnancy would have a significant impact on their locomotor abilities, resulting in an increase in wingbeat frequency and decrease in speed compared to non-pregnant bats, as well as delay achieving upward motion.

## MATERIALS AND METHODS

### Data collection

We filmed free-ranging big brown bats from a maternity colony at St. Patrick’s County Park (South Bend, Indiana, USA) on June 13 and July 22, 2021, representing seasonal periods of “pregnancy” versus “post-pregnancy”. The timing of birth for *E. fuscus* varies depending on latitude (Albrecht 2003; Boman 2018; Dood 1987; Krutzsch 1946; Lausen and Barclay 2006; Whitaker and Mumford 1971; Wisconsin Department of Natural Resources 2013; Workman 2008); we chose mid-June to ensure we recorded during late pregnancy based on a combination of both the published timing of peak *E. fuscus* birth closest to our location (late June, as reported in Dood 1987 for Northwestern Ohio) and five years of population monitoring at the site. The July date was selected to ensure most of the adults were in the post-pregnancy phase, but before most juveniles become volant. Emerging colony counts yielded 567 bats on June 9 and 887 bats on July 21, 2021. These counts indicate that at least 56% of the bats were pregnant in the June recording period, but this value was likely higher as pregnancy rates for *E. fuscus* generally exceed 90% (Kurta and Baker 1990) and we recorded before known peak juvenile volancy. Migration between roosts was assumed to be minimal; *E. fuscus* show high fidelity to undisturbed maternity roosts during the pregnancy period (Brigham and Fenton 1986), with dispersal typically beginning in August (Kurta and Baker 1990). Male *E. fuscus* do not live or roost with maternity colonies, so all adults were assumed to be female (O’Shea et al. 2021; Schmidly and Bradley 2016).

A high-resolution thermal imaging camera (FLIR a6751) was placed 1.6 meters above the ground and 15.6 meters from the barn, recording bats as they emerged from the main barn opening. Research followed ASM guidelines (Sikes et al. 2016). Most bats flew south and over a hedge located parallel to the barn door, so the camera field of view (FOV) was aligned starting at the edge of the barn and extending 10.4 meters along this flight line (Fig. 1) to record bats flying straight across the FOV. The camera was positioned in the same location between days, with alignment and position confirmed by aligning the FOV and comparing the pixel lengths of eight landmark features in the background of the videos.

**Fig. 1.**
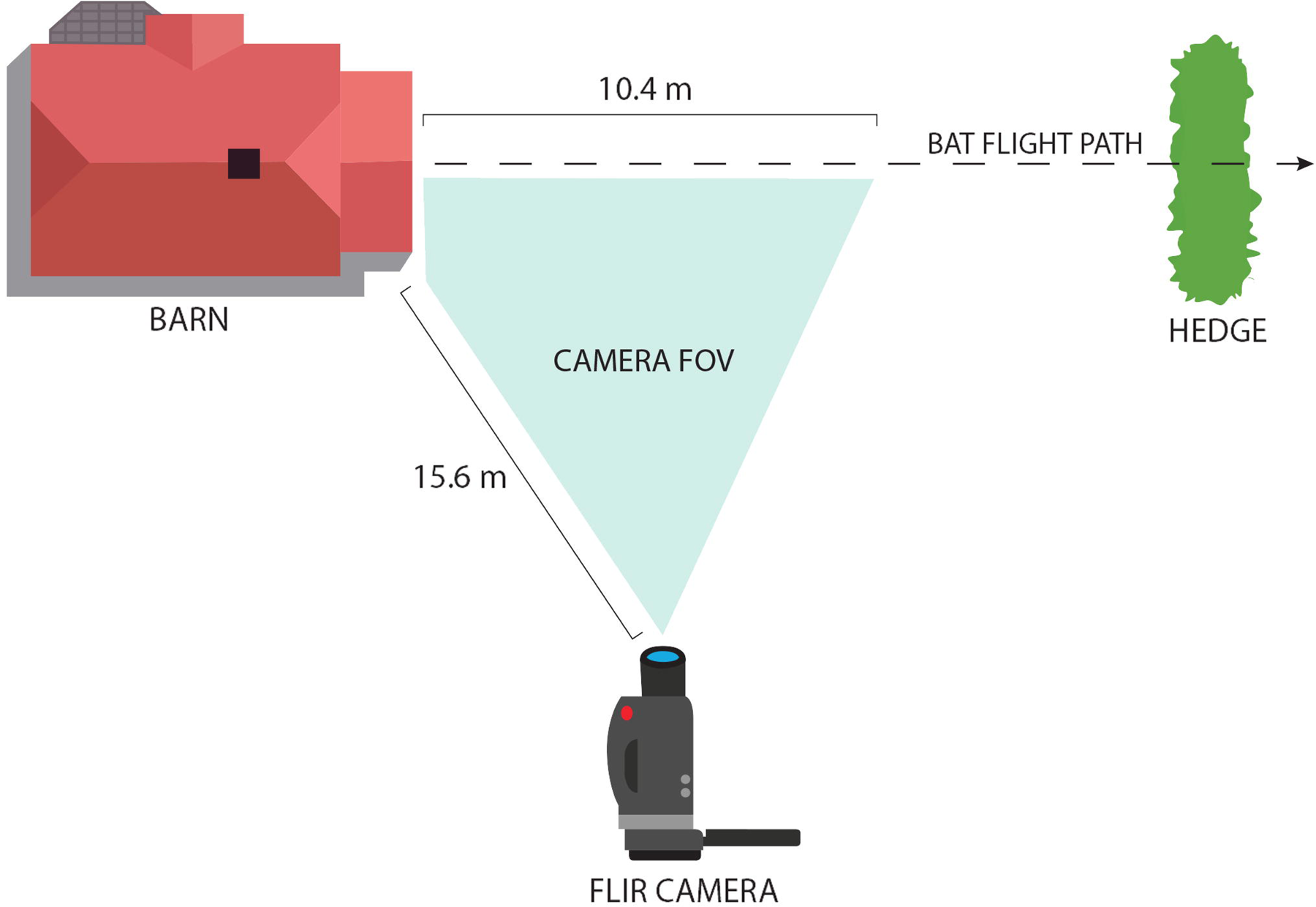
Plan view of camera setup and flight path of big brown bats (*Eptesicus fuscus*) recorded at St. Patrick’s County Park (South Bend, Indiana), 2021.

Video was captured at 125 Hz for the entirety of emergence. We used the following selection criteria for inclusion in analysis: the bat (1) did not emerge or fly in coordination with any other bats, (2) flew between a hedge parallel to the FOV, (3) engaged in flapping flight without coasting or swooping, and (4) visibly raised a wing before takeoff (as this was used as criterion for start of tracking). Because of the possibility that we recorded newly volant juvenile bats in July, we separated individuals based on the total body length: ≥ 116 mm adults, < 116mm juveniles, consistent with published body length measurements (Schmidly and Bradley 2016).

### Data analysis

Wingtip and center of body along the lower midline were tracked for each bat across frames using DLTdv8a (Hedrick 2008) (Fig. 2), starting with the first frame after emergence in which the wing visibly started to raise and continuing until the bat exited the FOV. The flight path was considered in two segments: before upward motion (UM) and after UM, with UM defined as the minimum vertical point along the curved flight path (Fig. 3).

**Fig. 2.**
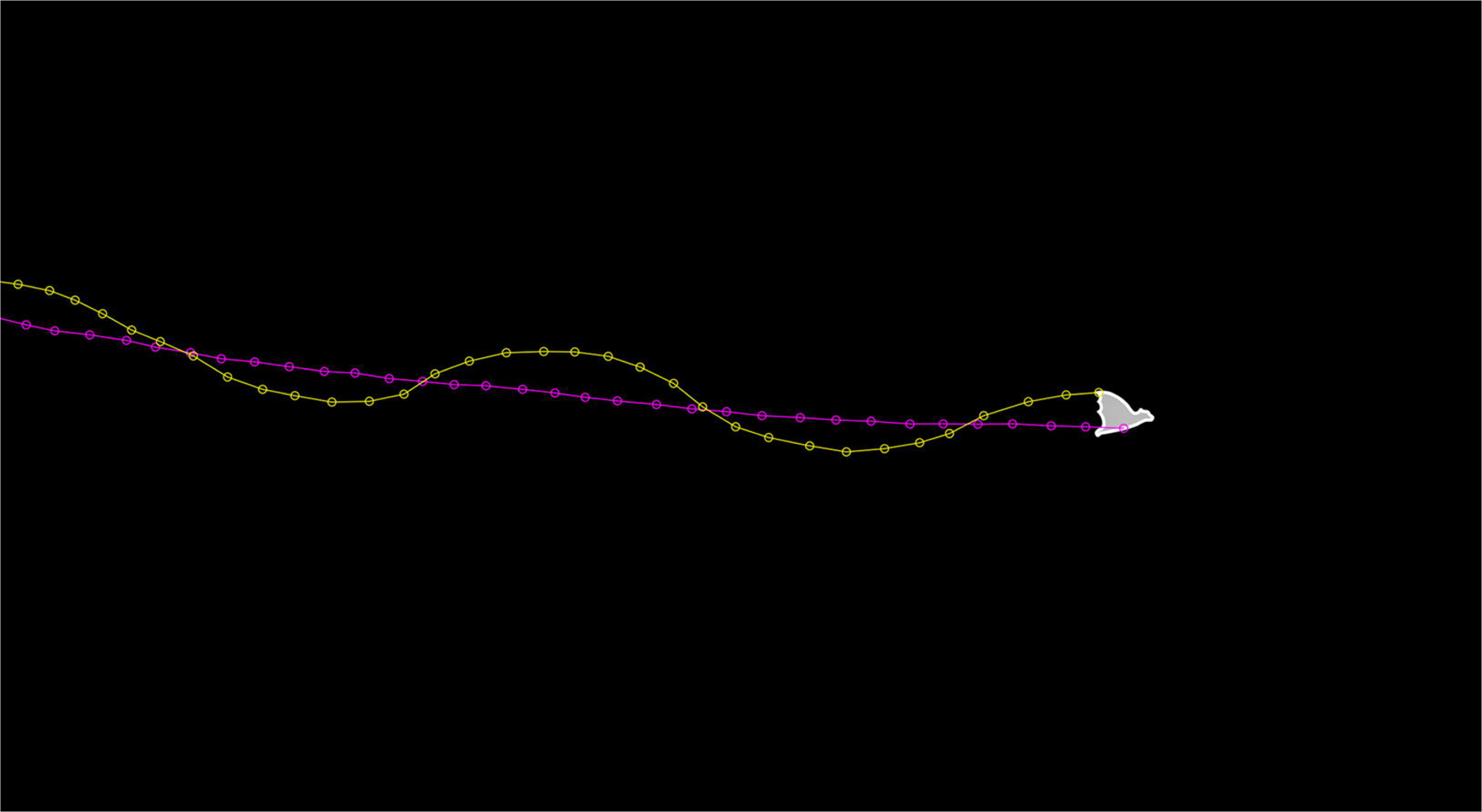
Example of bat tracked in DLTdv8a, showing wingtip (yellow) and center of body (purple).

**Fig. 3.**
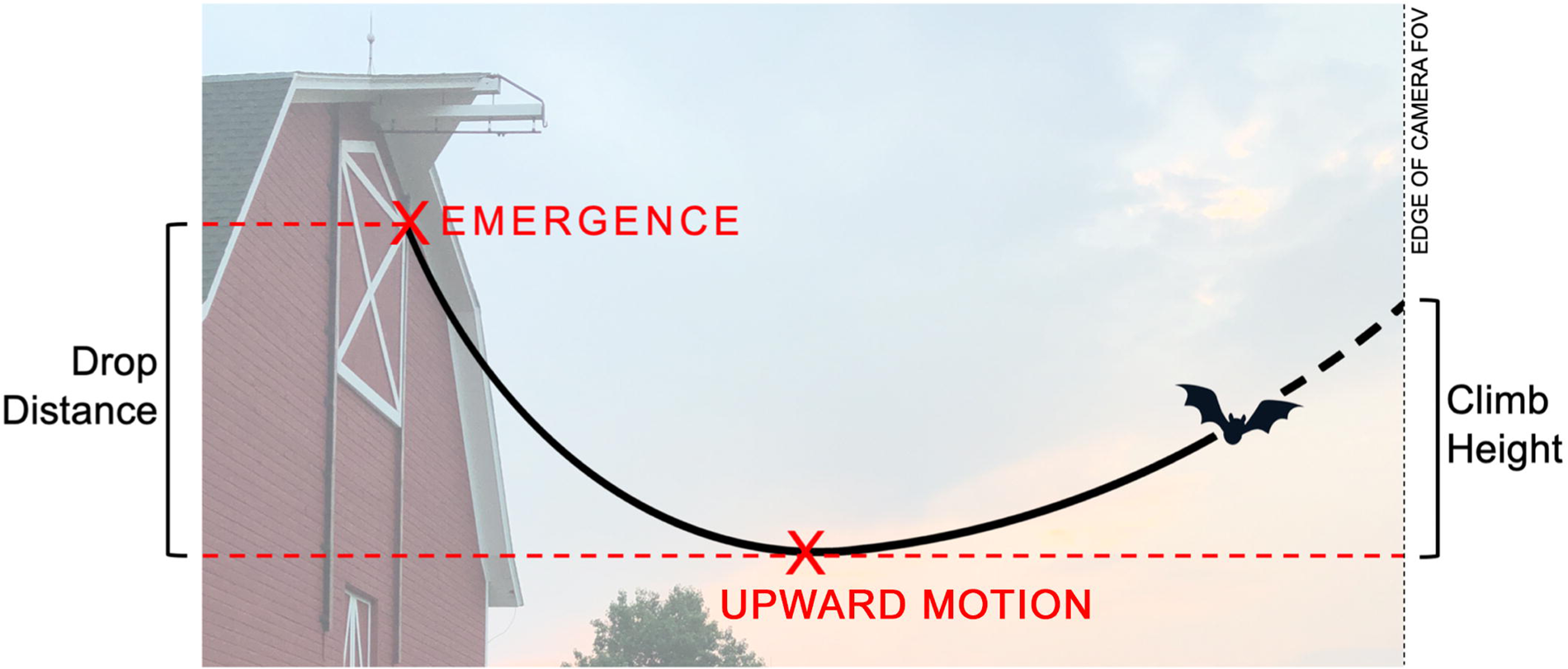
Illustration of key flight landmarks and variables of tracked bats.

From the tracking data and known camera frame rate, we calculated each bat’s drop distance, time from emergence until UM, average speeds before and after UM, climb height, and average wingbeat rate (WBR). Distances were converted to meters using a pixel-to-meter ratio as described in Kloepper and Bentley 2017. Average speed before UM was calculated as the distance (approximated as a straight line) between the center of body point at emergence and at UM, divided by the time difference between the frames. Average speed after UM was calculated similarly using the distance between the center of body at UM and the last tracked frame. All calculations were done in Matlab (R2021b).

We conducted a Shapiro-Wilk normality test on data for each variable of interest. Differences of means were analyzed using two-tailed independent samples t-test and Mann-Whitney *U* test. Interaction between variables was analyzed using ANCOVA. Statistical analysis was done in SPSS (v. 28.0.1.0) and Matlab (R2021b, MathWorks 2021).

## RESULTS

A total of 282 bats were recorded with the camera in June, and 394 were recorded in July. After applying our selection criteria, the final sample sizes for analysis were 28 bats in June “pregnancy period” and 24 bats in July “post-pregnancy period.” Application of the length criteria for the June bats resulted in 25 adults (126.83 ± 7.40 mm in length) and 3 juveniles (109.95 ± 5.21 mm), which we removed from further analysis; in July we recorded 11 adults (121.92 ± 2.25 mm) and 13 juveniles (108.97 ± 3.07 mm). There was no significant difference between July juveniles and July adults in drop distance, time to UM, speed before UM, climb height, or WBR (*p* > 0.05, Table 1); however, July juveniles had a significantly slower speed after UM (*t*(20) = 2.334, *p* = 0.030, Table 1), so to compare the June (pregnancy) versus July (post-pregnancy) period we considered only adults for further analysis. For speed after UM and climb height, we further removed 3 individuals that achieved UM 95% of the way through their measured flight path, as calculation of these parameters was not possible, resulting in a sample of 23 adults for June and 10 for July.

**Table 1.**
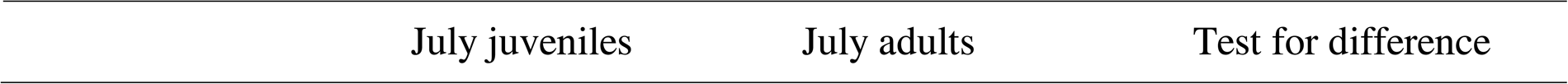
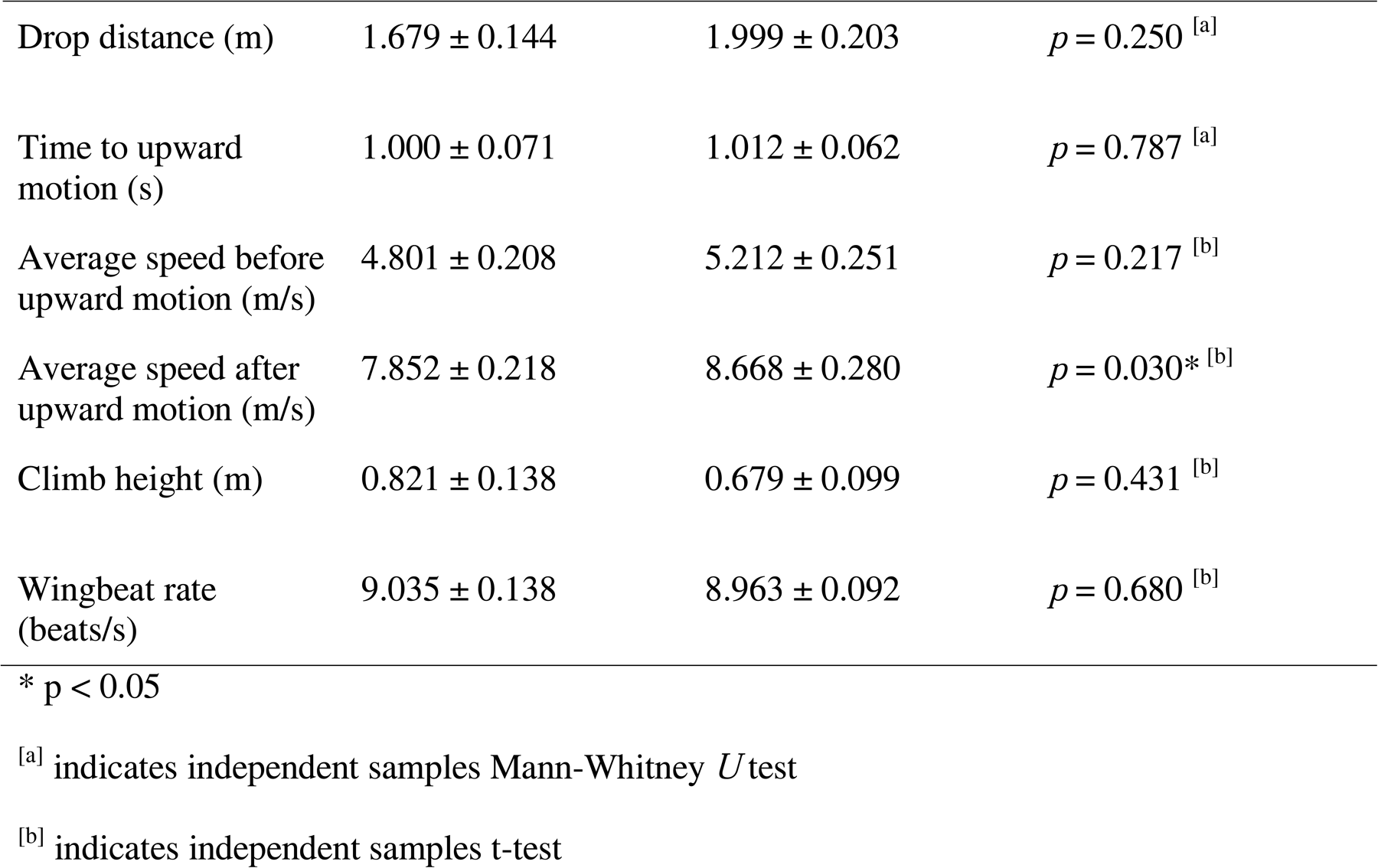
Mean ± SE for flight characteristics, as well as p-value from difference tests comparing July juvenile and July adult big brown bats (*Eptesicus fuscus*) recorded in South Bend, Indiana, 2021.

June adults had a significantly larger drop distance (Mann-Whitney *U* = 14.687, *p* = 0.004), longer time to UM (*t*(34) = 3.711, *p* < 0.001) and faster speed before UM (*t*(34) = 2.563, *p* = 0.015) compared to July adults (Table 2, Fig. 4A-C). Speed after UM was not significantly different between June and July adults (*t*(31) = 1.906, *p* = 0.066) (Table 2, Fig. 4D). June adults had a significantly shorter climb height and higher WBR than July adults (log transformation, *t*(31) = -2.921, *p* = 0.006; *t*(34) = 2.855, *p* = 0.007; Table 2, Fig. 4E-F). ANCOVA tests with drop distance as a covariate showed a marginally non-significant difference between WBR of June and July adults (*F*(1,33) = 4.112, *p* = 0.051); there was no significant difference for time to UM, speed before UM, speed after UM, or climb height (*p* > 0.05).

**Fig. 4.**
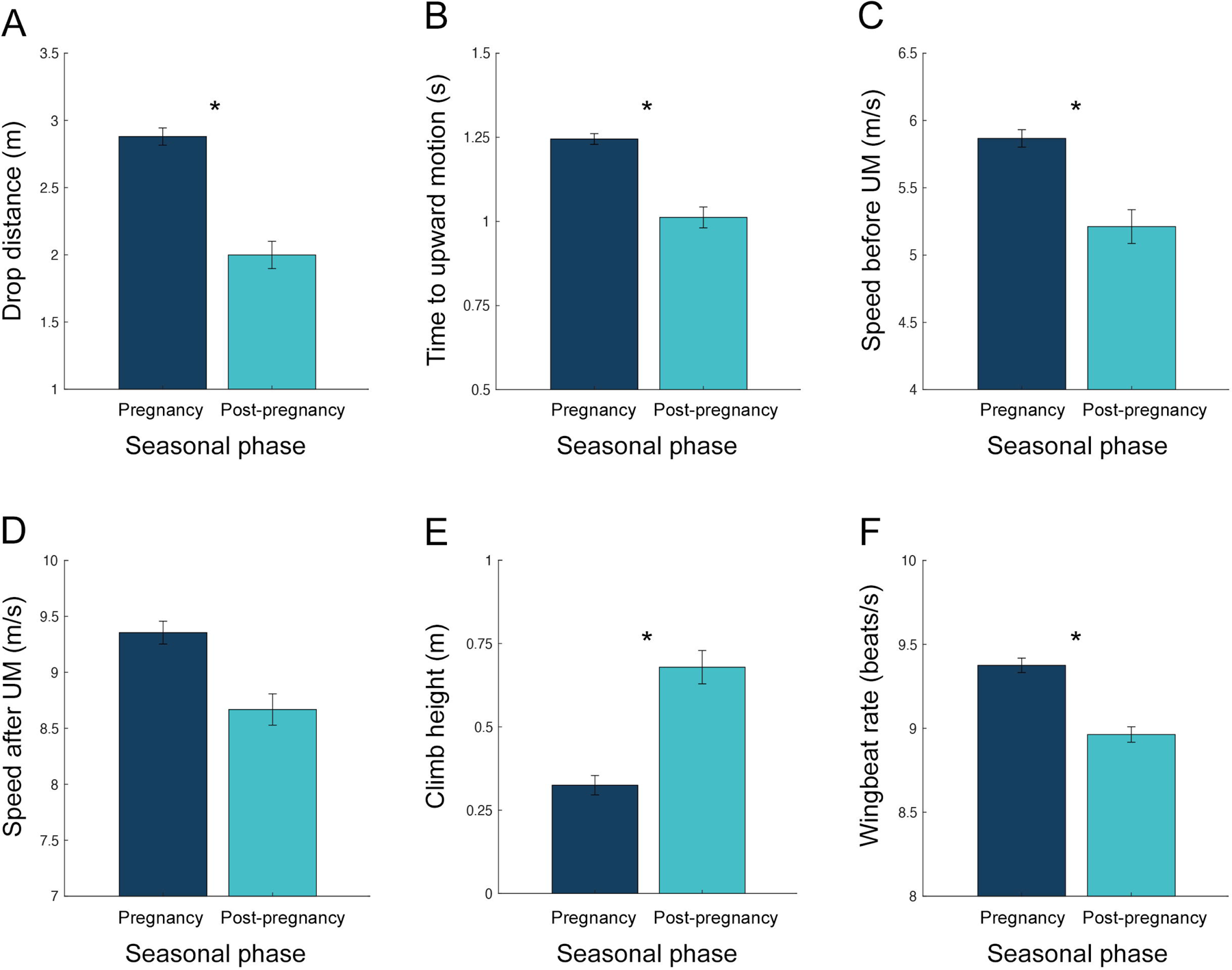
Comparing flight characteristics between June adult (navy) and July adult (light blue) big brown bats (*Eptesicus fuscus*) recorded in South Bend, Indiana, 2021. Bar graphs display (A) drop distance, (B) time to upward motion (UM), (C) speed before UM, (D) speed after UM, (E) climb height, and (F) wingbeat rate. Error bars indicate standard error. Asterisks indicate statistically significant difference.

**Table 2.** Mean ± SE for flight characteristics, as well as p-value from difference tests comparing June adult and July adult big brown bats (*Eptesicus fuscus*) recorded in South Bend, Indiana, 2021.

|  | June adults | July adults | Test for difference |
| --- | --- | --- | --- |
| Drop distance (m) | 2.880 ± 0.128 | 1.999 ± 0.203 | $p = 0.004^*$ <sup>[a]</sup> |
| Time to upward motion (s) | 1.245 ± 0.032 | 1.012 ± 0.062 | $p < 0.001^*$ <sup>[b]</sup> |
| Average speed before upward motion (m/s) | 5.867 ± 0.129 | 5.212 ± 0.251 | $p = 0.015^*$ <sup>[b]</sup> |
| Average speed after upward motion (m/s) | 9.355 ± 0.204 | 8.668 ± 0.280 | $p = 0.066$ <sup>[b]</sup> |
| Climb height (m) | 0.325 ± 0.058 | 0.679 ± 0.100 | $p = 0.006^*$ <sup>[c]</sup> |
| Wingbeat rate<br>(beats/s) | $9.375 \pm 0.086$ | $8.963 \pm 0.092$ | $p = 0.007^* \text{ }^{[b]}$ |

\* $p < 0.05$
<sup>[a]</sup> indicates independent samples Mann-Whitney $U$ test
<sup>[b]</sup> indicates independent samples t-test
<sup>[c]</sup> indicates independent samples t-test after log transform

## DISCUSSION

Our results suggest that pregnancy has a significant effect on flight in big brown bats, with a particularly strong impact on achieving UM after emergence. Adults recorded in the pregnancy phase had a significantly larger drop distance, longer time to UM, and shorter climb height compared to adults tracked in the non-pregnant phase (Table 2, Fig. 4-5). These differences suggest that pregnancy alters flight performance, increasing the distance and time a bat drops when beginning flight from a stationary point, and consequently requiring more power to achieve UM (Swartz et al. 2012). Non-pregnant bats were able to achieve UM and maintain an upward trajectory much more quickly than pregnant bats, likely because they did not need to adjust for a greater drop during the initial emergence phase. ANCOVA results indicate that differences in climb height and time to UM are likely the result of a greater drop distance experienced by pregnant bats, and not some inherent biomechanical difference due to pregnancy. Speed before UM was higher in pregnant bats, which may be attributed to acceleration over a longer time to UM, as well as higher wingbeat rate contributing thrust (Swartz et al. 2012). Juveniles and non-pregnant adults recorded in July showed similar flight characteristics (Table 1), suggesting that flight is well formed by the time of volancy.

**Fig. 5.**
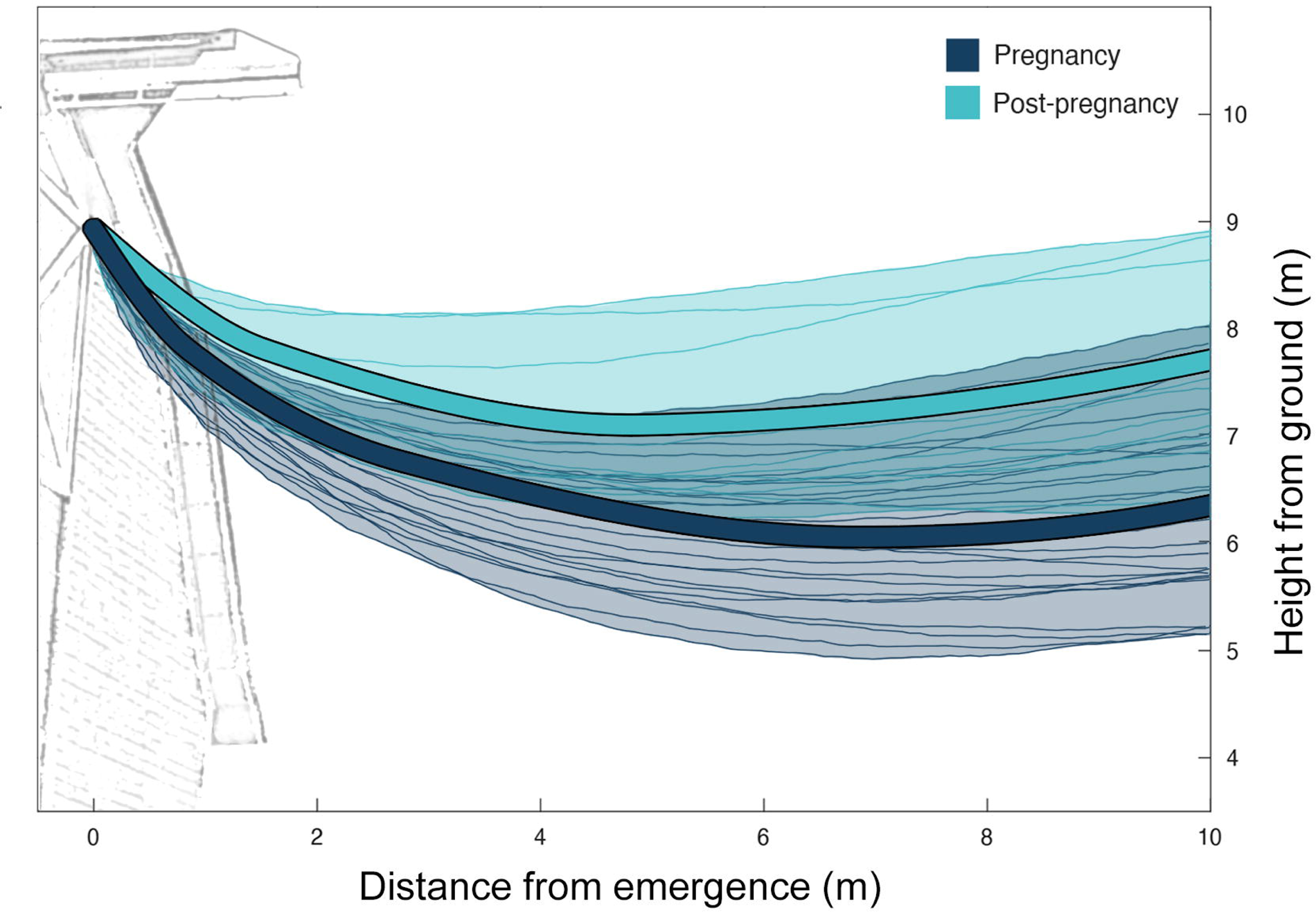
Trajectory plot showing individual flight paths and average trajectories of June adult (navy) and July adult (light blue) big brown bats (*Eptesicus fuscus*) recorded in South Bend, Indiana, 2021. Curve fits were generated using average coordinates of emergence, upward motion, and end points.

Wingbeat rate was significantly higher in pregnant bats, and this likely remains true even when controlling for drop distance at larger sample sizes than we were able to achieve. Given that the pregnant bats had more difficulty achieving and maintaining UM, it stands to reason that they would increase their wingbeat rate to generate the power required for flight (Fleuren 2017; Fokidis and Risch 2008; Hayssen and Kunz 1996; McLean and Speakman 2000; Swartz et al. 2012), which has been documented in prior work with bats undergoing horizontal, flapping flight (Hughes and Rayner 1991, 1993). This necessary power increase would be a regular, seasonal change that accompanies pregnancy-related increases in mass and wing loading, requiring a corresponding seasonal adjustment to flight behavior (Hughes and Rayner 1993).

Although there was no significant difference in speed after UM between the samples, pregnant bats tended to have faster speeds overall, which differs from prior results studying both the effect of mass (Hughes and Rayner 1991; Pennycuick et al. 1989; Videler et al. 1988) and pregnancy (Hughes and Rayner 1993) on flight speed across species. This likely resulted from greater potential energy converting to kinetic energy, as pregnant bats dropped over a greater drop distance. Additionally, pregnant bats likely experienced increased momentum while transitioning into UM, due to higher mass and higher flight speed before UM. In contrast, horizontal flight is almost entirely self-powered, and the force produced by the animals’ muscles needs to be greater in a heavier sample to compensate for the increased mass (Fokidis and Risch 2008; Hayssen and Kunz 1996), limiting the flight speeds they are able to achieve (Rayner 1999). If monitoring had continued further after UM, we predict trends in speed would have reversed between pregnant and non-pregnant bats as locomotion shifted to more closely resemble horizontal flight.

In our study, we assumed the adults sampled were pregnant or non-pregnant based on historical data for the species. Future work should validate this assumption by confirming pregnancy from capture data. Had we used larger samples with confirmed pregnant and non-pregnant adults, the differences we observed in flight characteristics may have been even more pronounced. Furthermore, pregnancy is a complex physiological process that involves many changes to an animal’s body and, consequently, locomotor abilities. In addition to increased body mass, locomotor performance could be affected by other pregnancy-related factors (e.g., morphological changes, energy reallocation/requirements) or seasonal environmental factors (e.g., temperature, humidity) (Kurta et al. 1990; Shine 2003). Future work could investigate the causal effect of increased body mass on flight performance in pregnant bats, controlling for these other factors. These important follow-up studies could help further reveal the impact that different pregnancy-related factors have on flight performance and the associated behavioral changes that bats exhibit in order to initiate and maintain UM during pregnancy.

## ACKNOWLEDGMENTS

We thank St. Patrick’s County Park for access to the site. We also thank Hannah Nichols and Valerie Eddington for assistance with data collection.

## DATA AVAILABILITY

Data available from Dryad Digital Repository.

## AUTHORS’ CONTRIBUTIONS

A.C.-Y.: conceptualization, data curation, formal analysis, investigation, methodology, writing— original draft, writing—review and editing; V.K H.Y.: conceptualization, investigation, methodology, writing—review and editing; M.K: resources, investigation, writing—review and editing; L.N.K.: conceptualization, formal analysis, funding acquisition, investigation, methodology, project administration, supervision, writing—review and editing.

All authors gave final approval for publication and agreed to be held accountable for the work performed therein.

